# A Mechanistic Model of Oncolytic Virus–CAR T Therapy Identifies Memory-Dependent Control of Solid Tumors

**DOI:** 10.64898/2025.12.24.696207

**Authors:** Neslihan N. Pelen, Trachette L. Jackson

**Affiliations:** Department of Mathematics and Statistics, University of South Florida, Tampa, USA; Department of Mathematics, Ondokuz Mayıs University, Samsun, Turkiye; Department of Mathematics, University of Michigan, Ann Arbor, USA

## Abstract

Chimeric antigen receptor (CAR) T-cell therapy can induce durable remissions in hematologic malignancies, yet its efficacy in solid tumors remains limited by antigen heterogeneity, an immunosuppressive tumor microenvironment, and poor T-cell persistence. Oncolytic viruses (OVs) provide a complementary therapeutic strategy by directly lysing tumor cells and reshaping local immune responses. Motivated by emerging evidence that OVs can also induce dual-specific, memory-like CAR T cells, we develop a unified mathematical framework that integrates these interacting modalities.

The model distinguishes antigen-positive and antigen-negative tumor populations, tracks key viral and immune compartments, and incorporates OV-mediated microenvironmental activation alongside CAR T-cell stimulation. This baseline formulation provides a platform for assessing whether additional mechanisms—such as OV-induced CAR T-cell memory—are necessary for durable tumor control. Sensitivity analysis of the memory-free system using partial rank correlation coefficients (PRCCs) identifies four dominant drivers of therapeutic outcome: antigen-negative tumor dynamics, viral kinetics, CAR T-cell cytotoxic effectiveness, and PD-1/PD-L1–mediated T-cell exhaustion. While these mechanisms can generate strong early responses, they are insufficient to prevent late relapse driven by antigen-negative tumor regrowth. Introducing a reduced memory variable to represent OV-driven CAR T recall fundamentally alters this behavior: enhanced memory induction and CAR T responsiveness sustain effector T-cell levels and enable long-term control of antigen-negative tumor populations. Together, these results highlight OV-induced CAR T-cell memory as a critical determinant of durable therapeutic efficacy in solid tumors.

## 1 Introduction

Chimeric antigen receptor (CAR) T-cell therapy has produced transformative clinical responses in several hematologic malignancies, where engineered T cells can induce deep and durable remissions [14, 15]. Extending this success to solid tumors, however, remains a major challenge. Tumor antigen heterogeneity, physical barriers to immune infiltration, and a profoundly immunosuppressive tumor microenvironment (TME) collectively limit CAR T trafficking, persistence, and cytotoxic efficacy [5, 7, 9, 26]. These constraints have motivated combination strategies aimed at reshaping the TME and broadening immune engagement.

Oncolytic viruses (OVs) offer a promising complementary approach. In addition to directly lysing tumor cells, OVs generate danger-associated molecular patterns, enhance antigen release, and induce cytokine and chemokine programs that recruit and activate innate and adaptive immune cells [3, 23, 33, 36]. Through these mechanisms, OVs can partially reprogram the TME to support immune infiltration, amplify antitumor immunity, and expand antigenic breadth [4, 36]. As a result, OV–CAR T combinations have emerged as a particularly attractive strategy for overcoming immune exclusion and dysfunction in solid tumors [20, 24, 29].

Immune checkpoint pathways further modulate these interactions. Following T-cell activation or viral infection, PD-1 and PD-L1 are rapidly upregulated, transmitting inhibitory signals that attenuate antitumor immune responses that would otherwise be effective [6, 16, 27, 32]. Checkpoint-mediated exhaustion therefore represents a critical barrier to both innate and adaptive immunity and remains a central focus of modern cancer immunotherapy [13, 19, 21, 22]. In combination settings, checkpoint signaling introduces additional feedback loops linking viral dynamics, immune activation, and CAR T persistence.

The resulting interplay among tumor heterogeneity, viral replication, innate and adaptive immune responses, CAR T activity, and checkpoint regulation generates tightly coupled, multiscale dynamics that are difficult to resolve experimentally. Mathematical modeling has therefore become an essential tool for dissecting how these processes jointly shape therapeutic efficacy, as demonstrated in prior studies of oncolytic virotherapy and CAR T treatment [12, 2, 17, 30, 31, 34]. For example, Storey et al. [30] showed that innate immunity can both support adaptive responses and prematurely eliminate the virus, with viral infectivity, innate clearance, and PD-1 signaling emerging as dominant determinants of tumor control. Separately, Kara et al. [17] demonstrated that antigen loss fundamentally limits CAR T efficacy in solid tumors and that increasing CAR T dose alone cannot overcome this barrier, highlighting the importance of tumor heterogeneity and cooperative killing mechanisms.

Recent experimental evidence suggests that OV–CAR T synergy may extend beyond TME remodeling and cytotoxicity. Evgin et al. [8] showed that OV infection can activate the native T-cell receptor (TCR) on CAR T cells via viral antigens, generating dual-specific CAR T cells capable of recognizing both tumor cells and virus-infected cells. These cells exhibit enhanced expansion, persistence, and memory-like properties, enabling long-term tumor control and recall responses upon viral re-exposure. These findings identify native-TCR stimulation and memory formation as critical determinants of durable efficacy in OV–CAR T combinations.

Motivated by these observations, we develop an integrated mechanistic model that unifies heterogeneous tumor populations, oncolytic viral dynamics, innate and adaptive immune responses, CAR T-mediated cytotoxicity, and PD-1/PD-L1–dependent regulation within a single mathematical framework. We first analyze a baseline formulation that excludes memory effects, using global sensitivity analysis to identify dominant biological axes governing treatment response. While this memory-free system captures strong early tumor reduction, it fails to prevent late relapse driven by antigen-negative tumor cells within biologically realistic parameter ranges. Guided by experimental evidence, we then introduce a reduced memory axis through which OV exposure promotes the emergence of a longer-lived, recall-capable CAR T population.

Through comparative simulations, we demonstrate that virus-driven CAR T memory fundamentally alters system dynamics, enabling sustained CAR T persistence and durable control of antigen-negative tumor populations. Together, these results identify OV-induced CAR T memory as a necessary mechanism for long-term efficacy in solid tumors and provide a unified quantitative framework for evaluating combination virotherapy–CAR T strategies.

In contrast to prior OV–CAR T models that focus on early cytotoxic synergy or check-point modulation, this work identifies immune memory as a structural requirement for durable control. By integrating tumor heterogeneity, viral dynamics, immune regulation, and memory-dependent CAR T persistence within a single framework, the model explains why enhanced early responses often fail and how virus-driven recall can fundamentally shift long-term outcomes.

## 2 Methods

To capture the coupled interactions among tumor cells, oncolytic viruses (OVs), CAR T cells, and the host immune system, we develop a mechanistic mathematical model that links viral infection, immune activation, and therapeutic response within a unified framework. The model is designed to reflect the key biological processes governing combination OV–CAR T therapy while remaining sufficiently structured to enable systematic analysis of treatment dynamics and sensitivity.

### 2.1 Mathematical Modeling Framework

The framework comprises four interacting components: (i) heterogeneous tumor populations, including antigen-positive and antigen-negative cells, each subdivided into susceptible and virus-infected states; (ii) free viral dynamics with repeated therapeutic dosing; (iii) innate and adaptive immune compartments, including natural killer (NK) cells, tumor-specific and virus-specific T cells, CAR T cells, and PD-1/PD-L1–mediated inhibitory signaling; and (iv) modulatory terms representing virus-driven reshaping of the tumor microenvironment (TME). Together, these components capture both the direct cytotoxic effects of therapy and the indirect immune-mediated mechanisms that shape long-term outcomes.

Our modeling assumptions are guided by experimental evidence that OV administration enhances CAR T-cell efficacy through two complementary mechanisms: inflammatory remodeling of the TME and direct stimulation of CAR T cells via their native T-cell receptor (TCR) [8]. We encode these effects using distinct viral modulation terms. First, virus-induced inflammatory activation of the TME is represented by a multiplicative activation function *S*(*V*), which amplifies innate and adaptive immune recruitment and activity in response to viral presence. This function is modeled as a saturating response to reflect biological limits on immune cell trafficking and cytokine-driven activation within the tumor.

Second, OV-driven stimulation of CAR T cells through native-TCR engagement is captured by a CAR T–specific modulation term *η*(*t, V*), which depends on both viral load and time. This term represents the emergence and recall of dual-specific CAR T subsets with enhanced expansion, persistence, and responsiveness upon repeated viral exposure. By explicitly separating TME-wide inflammatory activation from CAR T– intrinsic stimulation, the model allows these two biologically distinct pathways to be interrogated independently.

To simulate clinically relevant OV treatment regimens, viral administration is modeled as a sequence of finite-width dosing pulses implemented using a Gaussian pulse train [18]. These pulses act both as direct viral inputs and as temporal cues that synchronize immune activation and CAR T recall, enabling the study of treatment scheduling effects on therapeutic efficacy.

Under these assumptions, the model explicitly:

- resolves antigen-positive and antigen-negative tumor compartments, each divided into susceptible and virus-infected populations;
- tracks the principal viral and immune populations, including free virus *V*, NK cells *Z*, tumor-specific T cells *Y*_*T*_, virus-specific T cells *Y*_*V*_, CAR T cells *C*, and a PD-1/PD-L1–mediated inhibitory signal *P*;
- incorporates a virus-dependent TME activation multiplier *S*(*V*) that enhances innate and adaptive immune recruitment and function; and
- includes a virus- and time-dependent CAR T stimulation term *η*(*t*) representing dual-specific activation and memory-like recall.

This modeling approach, as shown in the schematic diagram in Figure 1, provides a biologically grounded yet analytically tractable platform for dissecting how viral dynamics, immune regulation, and tumor heterogeneity collectively determine the effect of OV–CAR T therapy in solid tumors.

**Figure 1.**
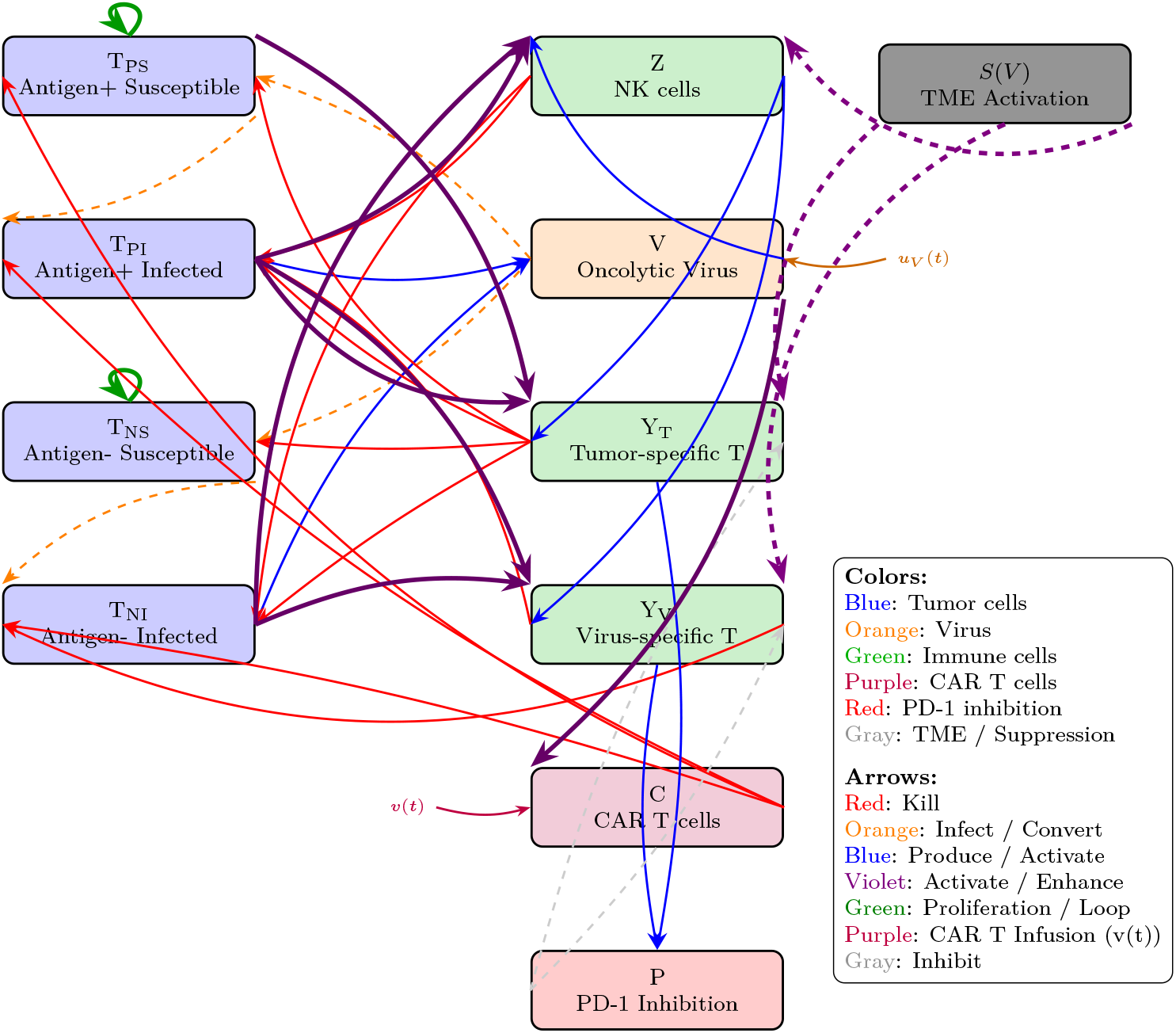
Interaction schematic for tumor, virus, immune effectors, CAR T, and PD-1 axis, including external viral input *u*_*V*_ (*t*) and CAR T infusion *v*(*t*).

### 2.2 Model equations

The governing equations, derived directly from the model schematic, are given below.

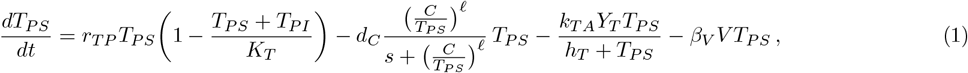

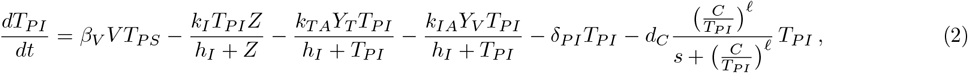

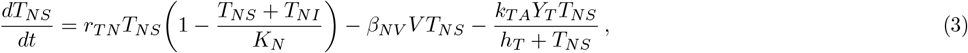

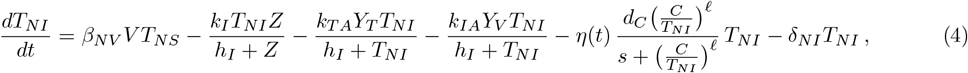

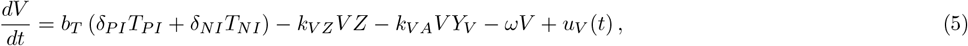

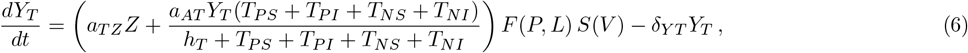

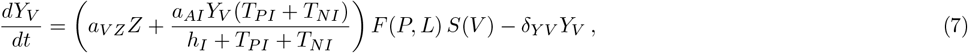

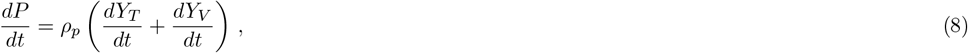

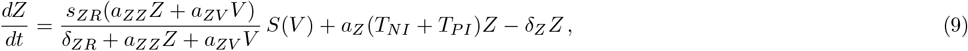

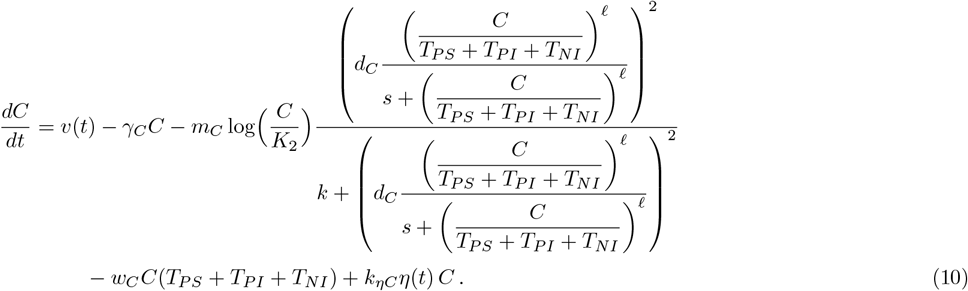

We collect here the scalar modulation functions used throughout the model. The PD-1/PD-L1–mediated suppression of effector *T* cells is modeled by

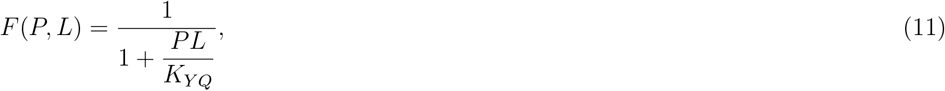

where the effective ligand level *L* is given by

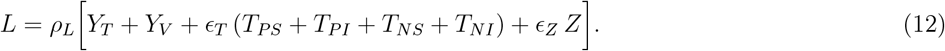

Virus-induced activation of the tumor microenvironment is captured by

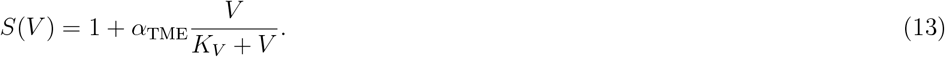

Dual-specific stimulation of CAR T cells by viral antigen is modeled through

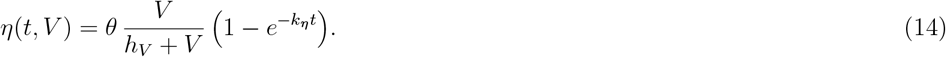

Finally, multiple OV injections are represented by the exogenous input

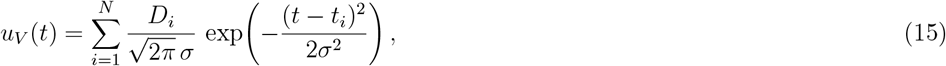

where (*t*_*i*_, *D*_*i*_) denote the injection times and doses, and *σ* = *V*_*σ*_ is the finite administration width [18].

The structure of the virus-induced activation *S*(*V*) and CAR T stimulation *η*(*t, V*) follows directly from experimental observations. For *S*(*V*), Evgin et al. [8] reported that OV replication drives inflammatory remodeling of the TME and enhances innate and adaptive immune recruitment. To represent this saturating immune activation, we used (13), where the constant term reflects basal immune tone and the saturating fraction captures the finite capacity of inflammation with increasing viral load.

For (14), the same study demonstrated that viral stimulation of the native TCR induces proliferation, memory formation, and long-term reactivation of dual-specific CAR T cells. To model this dynamic, *η*(*t, V*) was defined as a product of a viral load–dependent term 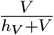 and a time-dependent build up 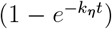, describing both viral sensitivity and progressive re-engagement following homologous boosting.

### 2.3 Memory-Extended Model

Motivated by the dual-specific memory phenotypes observed in Evgin et al. [8], we extend the baseline model by introducing a memory variable *M* (*t*) capturing the pool of dual-specific CAR T memory cells induced by viral antigen exposure. All baseline equations remain unchanged except for the CAR T stimulation term and the equations for *Y*_*T*_, *Y*_*V*_, *C*, and *T*_*NI*_, which are modified below, together with the additional memory state variable *M* (*t*).

#### 2.3.1 Memory variable and modified CAR T efficacy

We introduce a new dynamical variable *M* (*t*) obeying

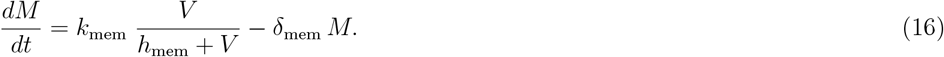

Here *k*_mem_ controls the rate at which viral antigen drives memory induction, *h*_mem_ is the viral load at which memory induction is half-maximal, and *δ*_mem_ captures slow contraction of the memory pool.

In the baseline (memory-free) model, dual-specific stimulation of CAR T cells is described by (14). To couple memory back to CAR T cells, we modulate this term as

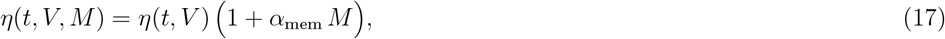

where *α*_mem_ quantifies the efficacy gain per unit memory. In addition, we include a small additive contribution +*σ*_*M*_ *M* to the activation of tumor-specific and virus-specific *T* cells, representing a general enhancement in responsiveness associated with a larger memory pool.

For convenience, denote *T*_tot_:= *T*_*PS*_ + *T*_*PI*_ + *T*_*NS*_ + *T*_*NI*_, *T*_Σ_:= *T*_*PS*_ + *T*_*PI*_ + *T*_*NI*_, and recall that *F* (*P, L*) and *S*(*V*) are given in (11) and (13). The tumor-specific *T* cells *Y*_*T*_ are now governed by

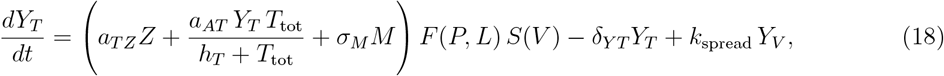

while the virus-specific *T* cells *Y*_*V*_ satisfy

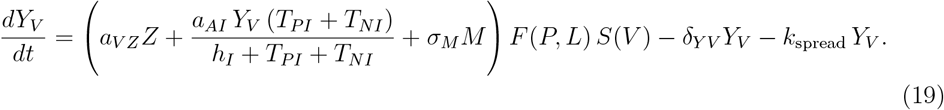

Here *k*_spread_ represents a slow, virus-induced drift from virus-specific to tumor-specific *T* cells, capturing epitope spreading.

The CAR T equation in the baseline model is (10), where the factor multiplying *m*_*C*_ is the saturation-type modulation term summarizing CAR T exhaustion. In the memory-extended model we replace the baseline stimulation term by the memory-amplified form in (17), yielding

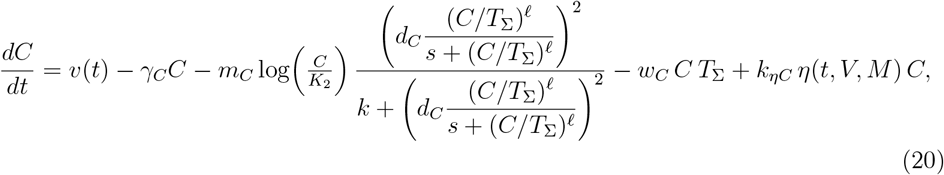

with *η*(*t, V, M*) given by (17).

Finally, in the tumor equation for antigen-negative infected cells (4), the CAR T killing term is also memory-modulated: the factor *η*(*t*) is replaced by *η*(*t, V, M*), i.e.,

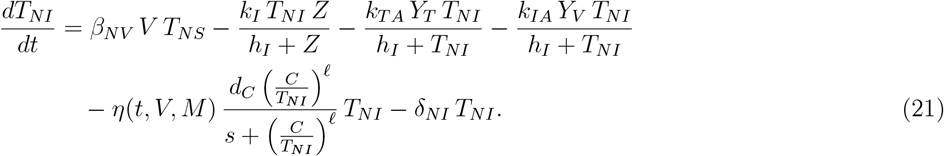

Equations (16)–(21) introduce a reduced memory/recall axis consistent with the qualitative trends reported in Evgin et al. [8]. Viral input *u*_*V*_ (*t*) increases the viral load *V* (*t*), which feeds into the memory variable *M* (*t*) through the saturating term *V/*(*h*_mem_ + *V*). The accumulated memory level persists beyond the dosing window and modulates CAR T stimulation via the factor (1 + *α*_mem_*M*) in *η*(*t, V, M*). The term *k*_spread_*Y*_*V*_ provides an additional mechanism for gradually shifting virus-specific activity toward tumor-directed activity.

### 2.4 Parameterization

The full list of model parameters, together with their biological interpretation and units, is summarized in Table 1 and 2.

**Table 1:** Model parameters, their biological interpretation, and units.

| Parameter | Description | Units |
| --- | --- | --- |
| $r_{TP}, r_{TN}$ | Growth rates of antigen positive and antigen negative tumor cells | $\text{day}^{-1}$ |
| $K_T, K_N$ | Carrying capacities of antigen positive and antigen negative tumor cells | $\text{mm}^3$ |
| $d_C$ | Maximum CAR T mediated killing rate | $\text{day}^{-1}$ |
| $\ell$ | Hill coefficient for CAR T killing | dimensionless |
| $s$ | CAR T killing half saturation constant | dimensionless |
| $k_{TA}$ | Killing rate coefficient of tumor specific T cells | $\text{day}^{-1}$ |
| $h_T, h_I$ | Half saturation constants for immune killing | $\text{mm}^3$ |
| $h_V$ | Viral load for half maximal CAR T activation | viral load<br>(model units) |
| $\beta_V, \beta_{NV}$ | Virus infection rates of antigen positive and antigen negative tumor cells | $(\text{viral load})^{-1} \text{day}^{-1}$ |
| $k_I, k_{IA}$ | Killing rates by NK cells and virus specific T cells | $\text{day}^{-1}$ |
| $\delta_{PI}, \delta_{NI}$ | Lysis rates of infected antigen positive and antigen negative tumor cells | $\text{day}^{-1}$ |
| $b_T$ | Viral burst per lysed tumor volume | viral load<br>(model units)<br>$\text{mm}^{-3}$ |
| $\theta$ | Strength of virus induced CAR T activation | dimensionless |
| $k_{VZ}, k_{VA}$ | Viral clearance rates by NK cells and virus specific T cells | $\text{mm}^{-3} \text{day}^{-1}$ |
| $\omega$ | Natural viral decay rate | $\text{day}^{-1}$ |
| $a_{TZ}, a_{VZ}$ | Activation of tumor specific and virus specific T cells by NK cells | $\text{day}^{-1}$ |
| $a_{AT}, a_{AI}$ | Autoproliferation of tumor specific and virus specific T cells | $\text{day}^{-1}$ |
| $\delta_{YT}, \delta_{YV}$ | Death rates of tumor specific and virus specific T cells | $\text{day}^{-1}$ |
| $\rho_p$ | Production coefficient for PD1/PDL1 regulatory signal | $\mu M \text{mm}^{-3}$ |
| $s_{ZR}$ | Baseline NK activation / source rate | $\text{mm}^3 \text{day}^{-1}$ |
| $a_{ZZ}$ | NK activation coefficient from NK cells | $\text{mm}^{-3}$ |
| $a_{ZV}$ | NK activation coefficient from viral load | $(\text{viral load})^{-1}$ |
| $\delta_{ZR}$ | NK activation saturation constant | dimensionless |
| $a_Z$ | Tumor induced NK proliferation coefficient | $\text{mm}^{-3} \text{day}^{-1}$ |
| $\delta_Z$ | NK cell death rate | $\text{day}^{-1}$ |
| $v(t)$ | CAR T infusion rate | $\text{mm}^3 \text{day}^{-1}$ |
| $\gamma_C$ | CAR T natural death rate | $\text{day}^{-1}$ |
| $m_C$ | CAR T exhaustion coefficient | $\text{mm}^3 \text{day}^{-1}$ |
| $K_2$ | CAR T carrying capacity in exhaustion term | $\text{mm}^3$ |
| $k$ | Saturation constant in exhaustion nonlinearity | $\text{day}^{-2}$ |
| $w_C$ | Tumor induced CAR T inhibition coefficient | $\text{mm}^{-3} \text{day}^{-1}$ |
| $K_{YQ}$ | PD1/PDL1 regulatory half saturation constant | $\mu M^2$ |
| $\eta(t, V)$ | Virus-induced CAR T activation factor | dimensionless |
| $\epsilon_T$ | Contribution of tumor cells to PD L1 expression | dimensionless |
| $\epsilon_Z$ | Contribution of NK cells to PD L1 expression | dimensionless |
| $\rho_L$ | Scaling factor for PD L1 expression | $\mu M \text{mm}^{-3}$ |
| $\alpha_{TME}$ | Strength of virus induced TME activation | dimensionless |
| $K_V$ | Saturation constant for virus-induced TME activation | viral load<br>(model units) |
| $k_{\eta C}$ | CAR T proliferation rate per unit $\eta(t)$ | $\text{day}^{-1}$ |

**Table 2:** Additional parameters introduced in the memory-extended model.

| Parameter | Description | Units |
| --- | --- | --- |
| $k_{\text{mem}}$ | Virus-driven memory induction rate | $\text{mm}^3 \text{ day}^{-1}$ |
| $h_{\text{mem}}$ | Half-saturation for memory induction | viral load (model units) |
| $\delta_{\text{mem}}$ | Memory decay rate (slow contraction) | $\text{day}^{-1}$ |
| $\alpha_{\text{mem}}$ | Memory efficacy multiplier | $\text{mm}^{-3}$ |
| $\sigma_M$ | Additive activation gain from memory | $\text{day}^{-1}$ |
| $k_{\text{spread}}$ | Epitope-spreading drift rate $Y_V \rightarrow Y_T$ | $\text{day}^{-1}$ |

#### 2.4.1 Parameters for Memory-free Model

All tumor and immune cell compartments *T*_*PS*_, *T*_*PI*_, *T*_*NS*_, *T*_*NI*_, *Y*_*T*_, *Y*_*V*_, *Z, C* are measured in mm^3^, time is measured in days, and *V* denotes viral load in model units. Parameter choices were informed by the biological ranges reported in prior OV–immunity and CAR T modeling studies [30, 17], while the specific numerical values were adapted to the structure and scaling of our model. The full parameter sets used in simulations are listed in Appendix A.

#### 2.4.2 Additional parameters for the memory-extended model

These parameters are introduced in Table 2 when simulating the memory-extended model in Section 2.3. The newly introduced variable *M* (*t*) represents the volume of dual-specific CAR T memory cells, which is measured in units of mm^3^. To analyze the effects of the memory-extended model, the parameter sets are provided in Appendix B.2.

## 3 Control Axes and Dynamical Outcomes in OV– CAR T Therapy

Across a wide range of parameter choices, the model exhibits consistent therapeutic outcomes driven by the coupled evolution of tumor growth, viral infection, and immune responses. Over time, this interaction produces distinct dynamical behaviors that shape tumor burden, viral persistence, and immune activity. Although strengthening viral replication and immune activation improves early tumor control, late relapse consistently occurs when immune memory is absent. This observation naturally leads to an extension of the model that incorporates immune memory.

### 3.1 Dominant Control Axes Identified by Global Sensitivity Analysis

We first identify the dominant biological axes governing tumor, viral, immune, and regulatory dynamics using global sensitivity analysis. The resulting sensitivity structure reveals four interacting control axes that organize the dynamics of the OV–CAR T–immune system.

Tumor growth and antigen-negative escape constitute the primary mechanism shaping long-term tumor burden. Sensitivity patterns indicate that late-time dynamics of the antigen-negative tumor population are dominated by intrinsic growth properties, reflecting tumor aggressiveness and carrying capacity rather than immune-mediated clearance. While viral infection and immune activation can transiently suppress tumor levels, their influence on antigen-negative cells weakens over time, allowing intrinsic proliferative dynamics to reassert control. As a result, long-term tumor burden is structured primarily by growth-driven escape mechanisms within the antigen-negative compartment (Fig. 2).

**Figure 2.**
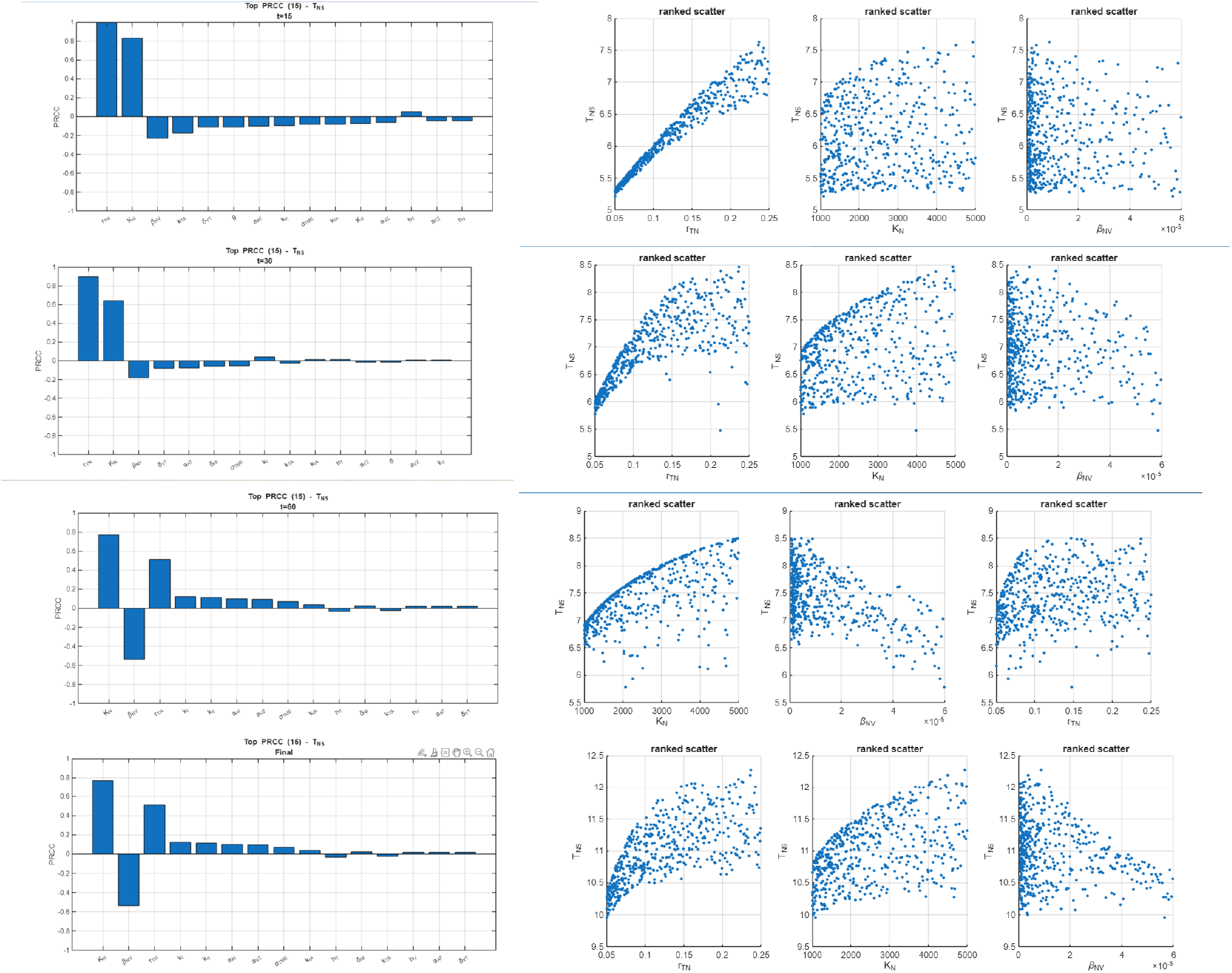
Global sensitivity for *T*_*NS*_ at *t* = 15, 30, 60, and final. Each row shows (left) the top PRCC magnitudes and (right) the ranked scatter plots for key parameters. Late-time antigen-negative tumor dynamics are governed primarily by intrinsic growth rather than immune-mediated suppression.

Viral dynamics are ultimately determined by the balance between infection-driven amplification and loss of antigen-negative infected tumor cells. At late times, viral persistence is governed by the ability of the virus to sustain a population of infected antigen-negative tumor cells, which requires continued infection of tumor cells and a sustained pool of susceptible antigen-negative cells supported by tumor growth and its carrying capacity. Earlier in the dynamics, rapid lysis of infected antigen-negative tumor cells can transiently suppress viral levels; however, this influence weakens over time if infection and tumor growth continue to replenish the infected compartment. When loss of infected antigen-negative tumor cells dominates, infected cells fail to accumulate, and viral replication cannot be sustained, leading to collapse of viral levels. These opposing effects define a shift in control over time and give rise to two distinct viral regimes: sustained infection supported by continued susceptibility of antigen-negative tumor cells, and viral elimination driven by rapid loss of virus-producing antigen-negative cells (Fig. 3).

**Figure 3.**
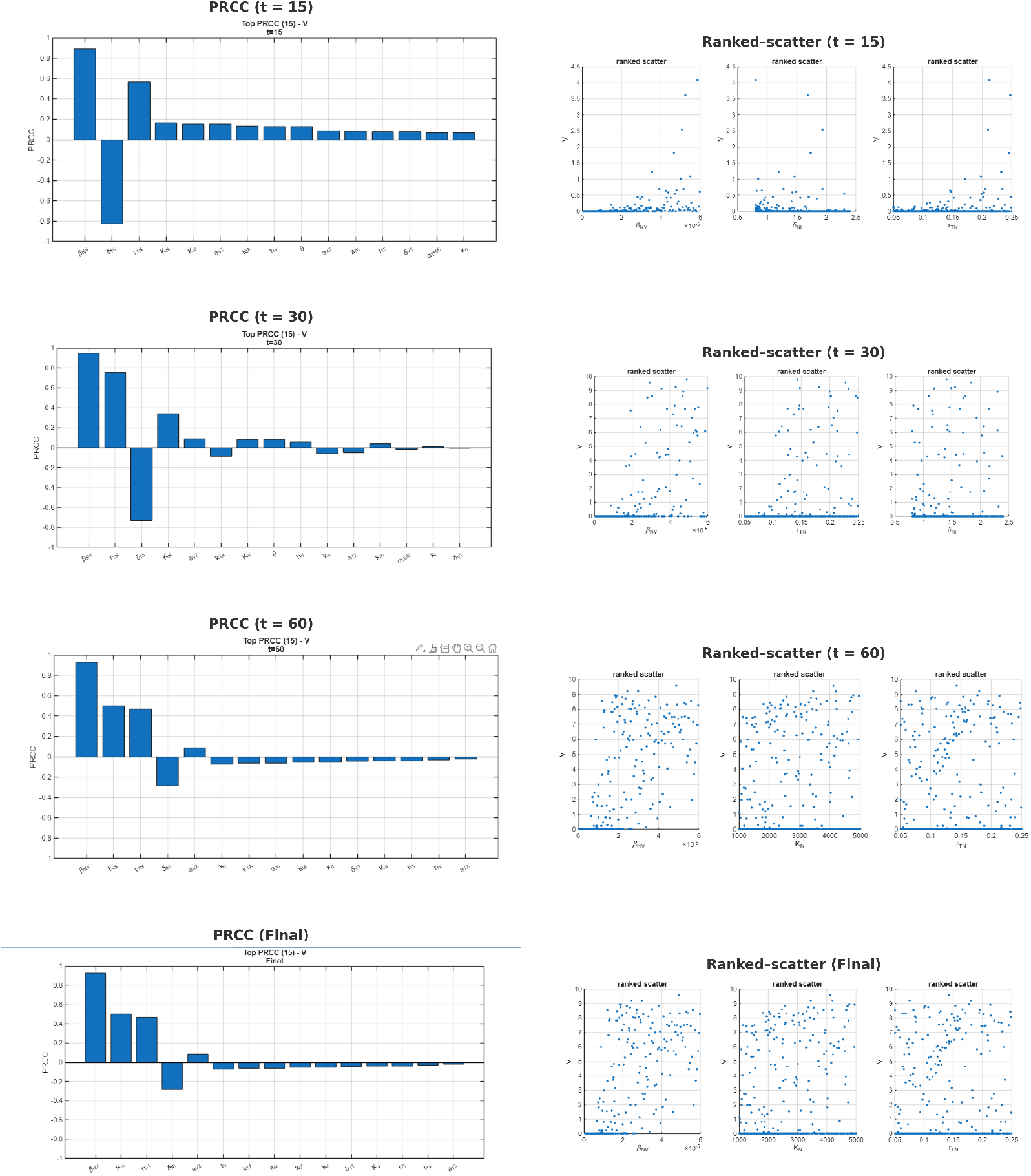
Global sensitivity for viral load *V* at *t* = 15, 30, 60, and final. Each row shows (left) the top PRCC magnitudes and (right) the ranked scatter plots for key parameters. Viral persistence is controlled by a balance between infection-driven amplification and loss of infected tumor cells, giving rise to sustained-infection or viral-clearance regimes.

CAR T dynamics follow an activation–persistence trade-off that evolves over time. Sensitivity patterns indicate that early CAR T expansion is governed primarily by intrinsic activation and cytotoxic efficacy, reflecting parameters that regulate CAR T responsiveness and tumor-directed killing. By contrast, sustained CAR T presence at later times becomes increasingly dependent on viral stimulation. As viral stimulation decreases, CAR T levels also decrease, even when intrinsic activation is strong, showing that long-term CAR T persistence becomes virus-dependent. This temporal separation explains why transient immune amplification alone fails to support long-term CAR T persistence in memory-free settings (Fig. 4).

**Figure 4.**
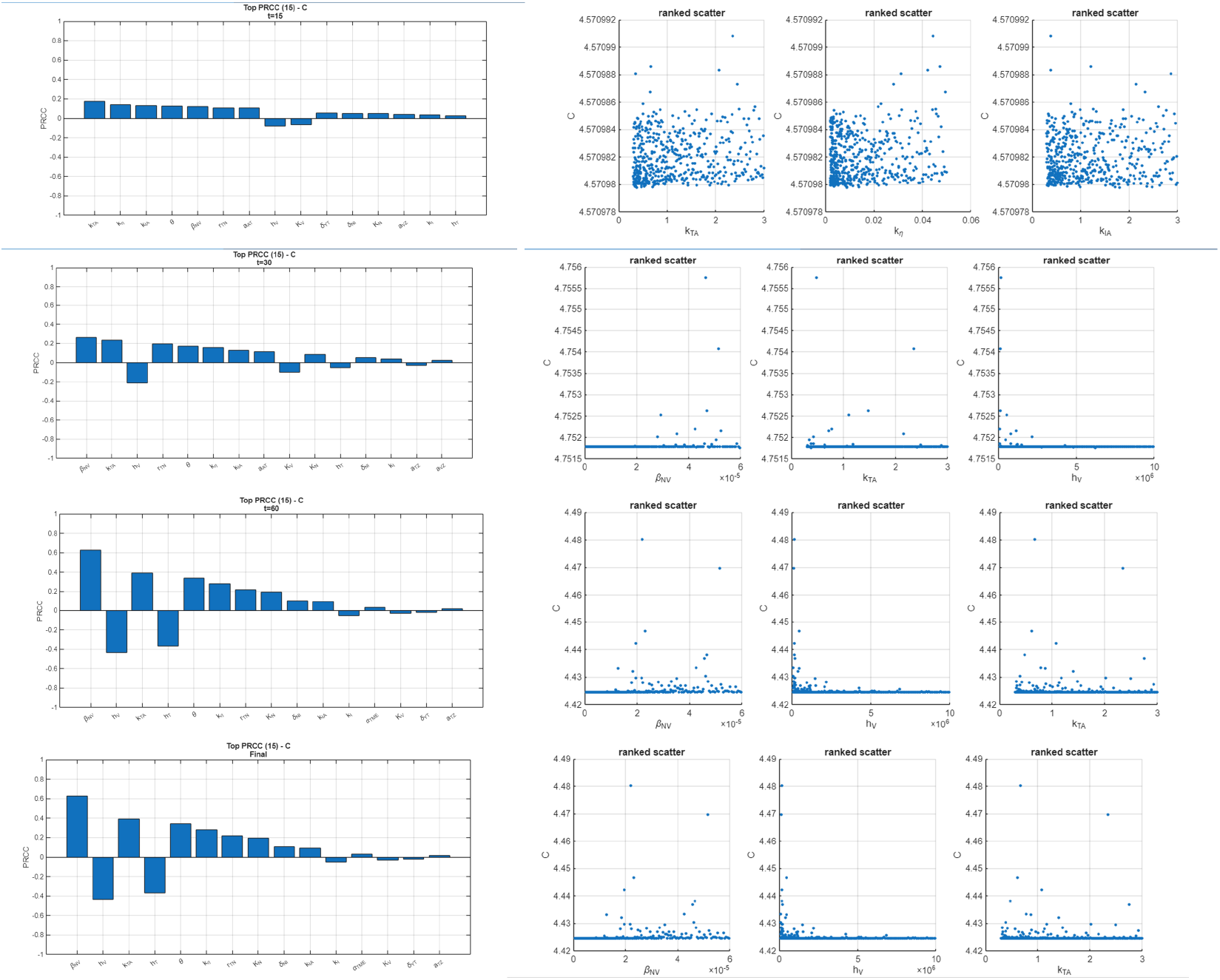
Global sensitivity for CAR T cells *C* at *t* = 15, 30, 60, and final. Each row shows (left) the top PRCC magnitudes and (right) the ranked–scatter for key parameters. Early CAR T expansion is activation-driven, whereas long-term CAR T persistence becomes dependent on sustained viral stimulation.

Immune exhaustion mediated through PD-1/PD-L1 signaling acts as a regulatory constraint that limits, but does not independently determine, therapeutic outcome. Sensitivity analysis indicates that PD-1/PD-L1 levels are tightly linked to the survival of tumor-specific effector T cells, serving as a marker of immune persistence and activation. In particular, higher death rates of tumor-killing T cells are associated with reduced PD-1/PD-L1 signaling, reflecting immune silencing due to effector cell loss rather than effective immune regulation. Conversely, strong immune activation arising from virus-specific T-cell cytotoxicity and NK-mediated activation of virus and tumor-specific T cells, together with viral infection of antigen-negative tumor cells and the resulting antigen-driven stimulation of CAR T cells, can transiently elevate PD-1/PD-L1 signaling by sustaining active tumor-killing effector populations. This elevation reflects ongoing immune engagement rather than immune failure. This axis therefore modulates and constrains the effectiveness of tumor and viral control, but does not independently determine the dominant growth- and infection-driven dynamics (Fig. 5).

**Figure 5.**
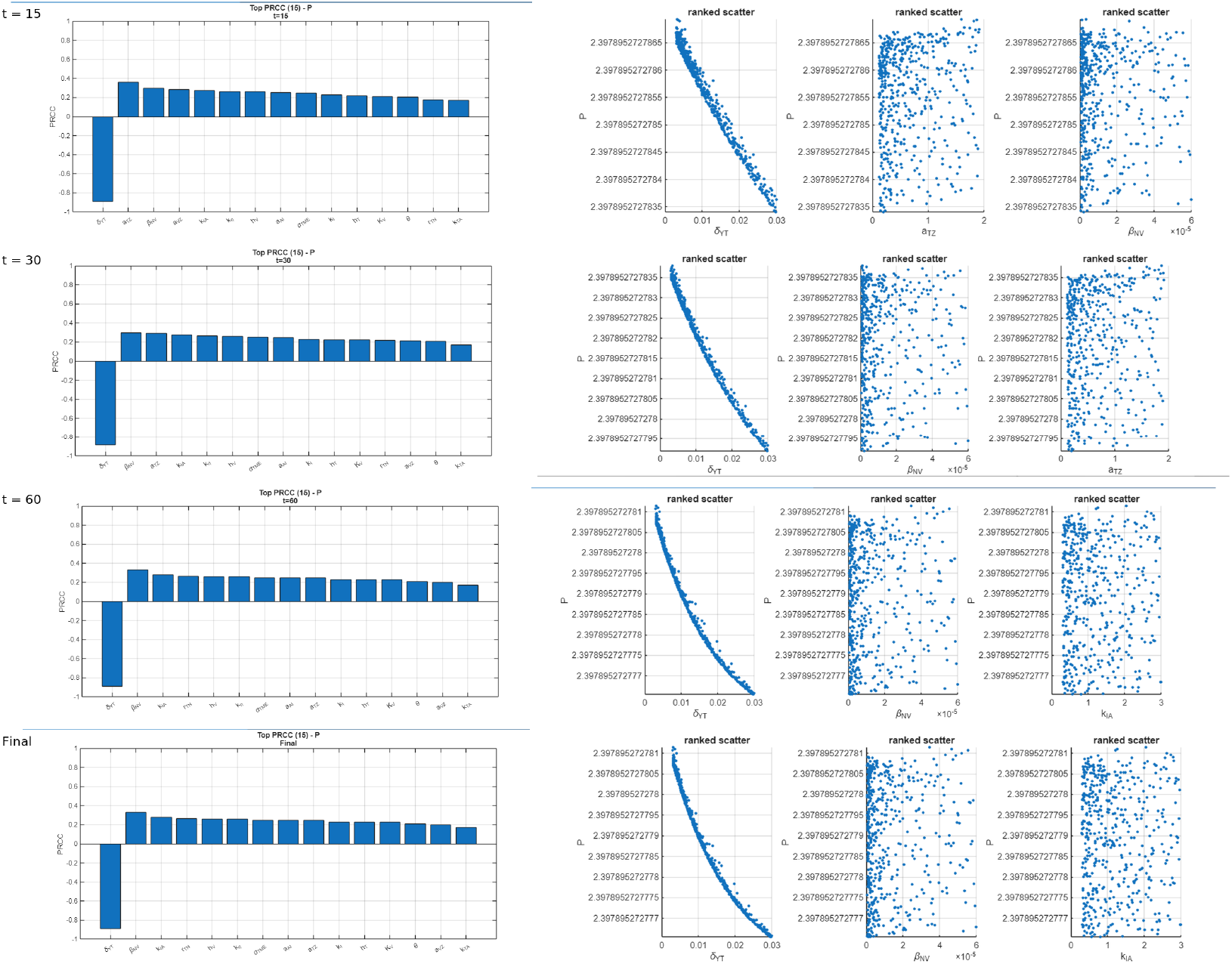
Global sensitivity for PD-1/PD-L1 level *P* at *t* = 15, 30, 60, and final. Across all times, *P* is dominantly and negatively associated with the tumor-specific T-cell death rate *δ*_*Y T*_, while infection/activation knobs (*β*_*NV*_, *k*_*IA*_, *a*_*T Z*_, and the CAR T efficacy terms *θ, k*_*η*_, *h*_*V*_) show mild positive influence. PD-1/PD-L1 signaling reflects ongoing immune engagement and constrains response efficiency, but does not independently determine therapeutic outcome.

Together, the PRCC results reveal four coupled control axes that structure all subsequent system behavior.

### 3.2 Sensitivity Structure of Tumor, Virus, CAR T, and PD-1/PD-L1 Dynamics

The dominant control axes identified above do not act in isolation. Instead, their interaction produces coordinated system-level behavior across tumor, viral, and immune compartments. In this subsection, we synthesize the sensitivity structure across variables to clarify how these axes shape qualitative dynamical regimes, rather than repeating parameter-level rankings.

#### 3.2.1 Antigen-Negative Tumor Dynamics

##### Result

Long-term antigen-negative tumor burden is governed primarily by intrinsic tumor growth.

Although viral infection and immune activation can transiently suppress tumor burden, late-time dynamics are dominated by the intrinsic growth of antigen-negative tumor cells. Sensitivity structure consistently indicates that tumor aggressiveness and carrying capacity determine long-term outcomes, creating a separation between early immune-driven responses and late growth-dominated behavior. This explains why enhanced immune activity alone cannot prevent antigen-negative tumor escape in memory-free settings.

#### 3.2.2 Viral Persistence versus Clearance Regimes

##### Result

Viral dynamics exhibit a threshold-like transition between sustained infection and elimination.

Sensitivity patterns reveal two qualitatively distinct viral regimes. When infection processes dominate, viral levels are maintained through continued infection of susceptible tumor cells. In contrast, when loss of infected cells dominates, virus-producing compartments fail to accumulate and viral replication collapses. Viral dynamics, therefore, serve as a central determinant linking tumor susceptibility to downstream immune sustainability.

#### 3.2.3 CAR T Expansion Is Virus-Dependent at Late Times

##### Result

Sustained CAR T levels at late times require ongoing viral stimulation.

Early CAR T expansion is driven primarily by intrinsic activation and cytotoxic efficacy. However, the sensitivity structure shows that CAR T persistence at later times becomes increasingly coupled to viral activity. As viral stimulation decreases, CAR T levels decline even when intrinsic activation parameters are strong. This shift from activation-driven to virus-coupled control explains why transient immune amplification alone cannot support durable CAR T persistence in memory-free scenarios.

#### 3.2.4 PD-1/PD-L1 as a Secondary but Constraining Axis

##### Result

Immune exhaustion modulates therapeutic effectiveness without independently determining outcome.

PD-1/PD-L1 dynamics are closely linked to the survival and activation of tumor-killing effector T cells. Elevated inhibitory signaling reflects ongoing immune engagement rather than autonomous suppression of tumor control. While immune exhaustion constrains response efficiency, the sensitivity structure indicates that it does not override the dominant growth- and infection-driven axes. Instead, this regulatory pathway shapes immune effectiveness within limits imposed by tumor proliferation and viral persistence.

Together, these synthesized sensitivity structures clarify how tumor growth, viral persistence, CAR T sustainability, and immune regulation jointly shape the system-level behaviors examined in the following dynamical analyses.

### 3.3 PRCC-Guided Construction of Aggressive and Enhanced-Activation Regimes

Based on the global sensitivity structure identified in Sections 3.1 and 3.2, we constructed two parameter regimes that selectively amplify the dominant biological axes. Specifically, these regimes strengthen the axes governing (i) the balance between tumor growth and infection-driven suppression and (ii) immune activation and persistence, which emerged consistently as the primary determinants of system behavior in the PRCC analysis.

#### *AGG*: aggressive baseline regime

The ***AGG*** regime applies *moderate shifts* along the PRCC-identified axes. Tumor–infection balance and immune activation are enhanced within biologically plausible ranges, yielding an aggressive but still conservative therapeutic configuration that reflects strengthened treatment effects without fundamentally altering the system structure.

#### *AGG+*: enhanced-activation regime

The ***AGG+*** regime pushes the *same control axes further* by amplifying viral output and increasing CAR T responsiveness to virus-associated stimulation. Compared to *AGG*, this regime represents a high-activation setting in which infection-driven exposure and immune activation are intensified, while remaining aligned with the sensitivity-guided mechanisms identified by PRCC.

Minor adjustments to the viral burst size *b*_*T*_ and decay rate *ω* were included to rescale viral exposure while remaining consistent with the PRCC-identified control axes.

Figure 6 illustrates how these PRCC-guided regime constructions modify the coupled tumor–virus–immune dynamics by selectively amplifying the dominant control axes. The figure provides the reference trajectories used to compare baseline, *AGG*, and *AGG+* behaviors in the dynamical analyses that follow. Complete parameter values and biological plausibility considerations for both regimes are provided in Appendix A.

**Figure 6.**
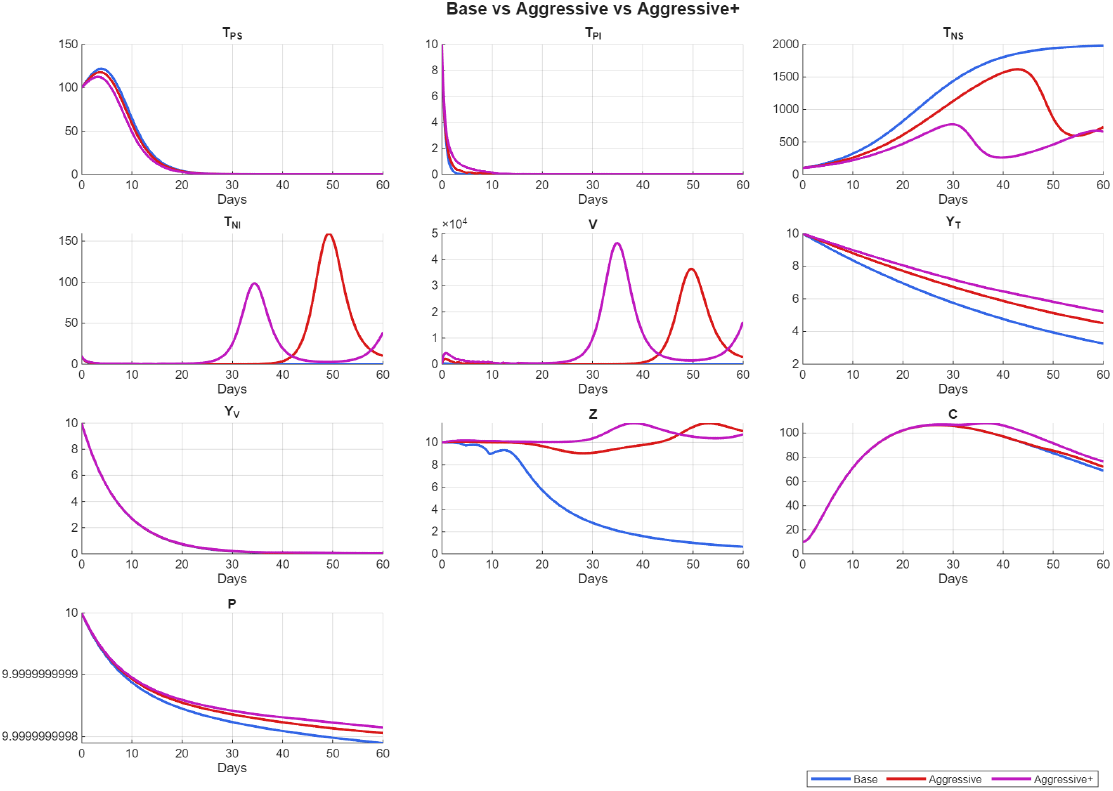
PRCC-guided construction of aggressive therapeutic regimes. Baseline, *AGG*, and *AGG+* simulations demonstrate how selective amplification of PRCC-identified control axes alters tumor burden, viral persistence, and immune activity without introducing new mechanisms.

### 3.4 Dynamical Outcomes Without Memory: Improved Early Control but Inevitable Relapse

#### Result

Despite strengthened viral replication and CAR T activation, all memory-free scenarios eventually relapse due to antigen-negative tumor regrowth.

Across baseline, *AGG*, and *AGG+* regimes, strengthening the sensitivity-identified mechanisms consistently improves early tumor suppression. In all cases, tumor burden is initially reduced through enhanced viral infection of the tumor cells and increased CAR T-mediated cytotoxicity, reflecting successful short-term amplification of effector mechanisms.

This early tumor suppression is accompanied by a transient expansion of CAR T cells. Viral infection and antigen exposure drive rapid CAR T activation and proliferation, producing a pronounced early peak in CAR T abundance. However, in the absence of immune memory, this expansion cannot be sustained once viral stimulation declines, leading to contraction of the effector population.

At later times, the loss of sustained immune pressure allows antigen-negative tumor cells to escape control progressively. The antigen-negative compartment *T*_*NS*_ re-emerges and ultimately dominates tumor burden, even under the highly activated AGG+ regime in which viral output and CAR T responsiveness are maximally enhanced within biologically plausible ranges. This late resurgence of *T*_*NS*_ occurs because intrinsic tumor growth becomes dominant once early effector mechanisms decrease.

Crucially, these outcomes demonstrate the failure of two intuitive intervention strategies in memory-free settings. Increasing CAR T killing alone does not stop relapse because CAR T cells decrease when viral stimulation decreases. Similarly, extending viral persistence alone fails to maintain long-term tumor control, since sustained viral presence cannot compensate for the absence of immune recall. In both cases, early effector amplification is transient and cannot be reinitiated after initial decline.

Together, these findings indicate that relapse arises from a structural limitation of memory-free dynamics rather than suboptimal parameter choices. Enhancing short-term effector activity is therefore insufficient for durable tumor control, motivating the inclusion of an explicit immune memory mechanism in the model.

#### Synthesis of Memory-Free Results and Implications

Up to this point, the analysis establishes several central results of memory-free OV–CAR T dynamics. First, system behavior is organized by a small number of interacting biological control axes rather than by fine-scale parameter tuning. These axes include intrinsic tumor aggressiveness and antigen-negative growth; the balance between viral infection and lysis; CAR T responsiveness to virus-associated stimulation; and PD-1/PD-L1-mediated immune exhaustion.

Second, CAR T dynamics exhibit a fundamental temporal shift in control. Early CAR T expansion is governed primarily by intrinsic activation and cytotoxic efficacy, whereas long-term CAR T persistence becomes increasingly dependent on sustained viral stimulation. In this late phase, the viral infection rate *β*_*NV*_ emerges as a dominant driver, linking CAR T maintenance to continued antigenic and inflammatory cues rather than to initial effector strength.

Third, and most critically, strengthening these control axes within biologically plausible ranges is insufficient to prevent late antigen-negative tumor escape in the absence of immune memory. Notably, even the *AGG+* regime—which maximizes viral output and CAR T responsiveness within realistic bounds—fails to prevent relapse, underscoring that this limitation is structural rather than parametric. In these settings, relapse arises not because parameters are poorly chosen, but because transient effector amplification cannot be re-engaged once immune pressure wanes.

Taken together, these results demonstrate that durable tumor control cannot be achieved in the memory-free system. Instead, sustained suppression of antigen-negative tumor populations requires a mechanism for long-term immune recall, motivating the explicit incorporation of immune memory in the following section.

### 3.5 Effect of Memory on Long-Term Tumor Control

Introducing a memory mechanism fundamentally alters late-time tumor dynamics, enabling durable suppression of antigen-negative tumor cells. While all treatment settings achieve rapid early reduction of antigen-positive tumor burden, their long-term outcomes diverge sharply once memory effects are considered.

Figure 7 compares the *Base, Aggressive, Aggressive+*, and *Cure-tuned++* parameter regimes under the memory-extended formulation.

**Figure 7.**
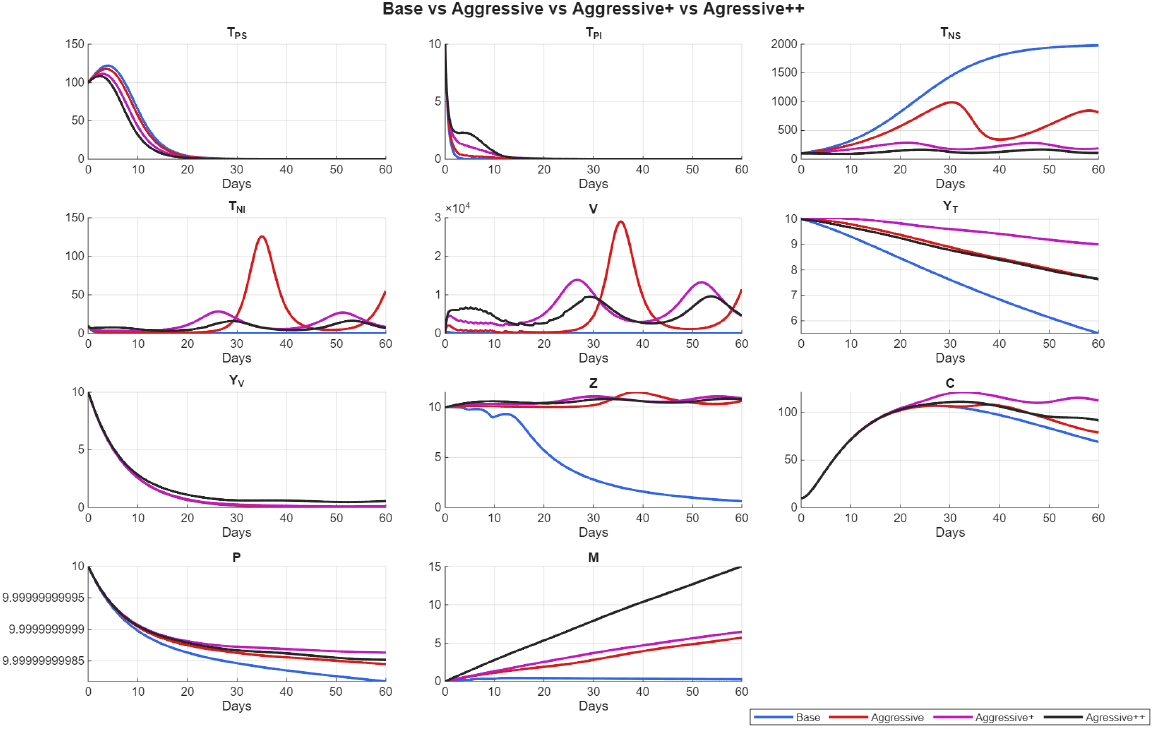
Comparison of tumor, immune, viral, regulatory, and CAR T dynamics under the four parameter regimes, ***Base, Aggressive, Aggressive+***, and ***Cure-tuned++***, in the memory-extended model. Immune memory stabilizes CAR T persistence at late times, converting transient early tumor control into durable suppression of antigen-negative tumor populations.

In all cases, antigen-positive tumor cells decline early, reflecting the immediate cytotoxic impact of OV infection and CAR T activity. However, in the *Base* regime, antigen-negative tumor cells gradually expand at later times, leading to tumor regrowth despite an initially favorable response.

The *Aggressive* regime produces a stronger early effect but induces a pronounced infection-driven wave around days 30–40, resulting in large transient fluctuations across tumor, viral, and immune compartments. In contrast, the *Aggressive+* and *Cure-tuned++* regimes exhibit more balanced dynamics. In these memory-enabled settings, antigen-negative tumor populations remain suppressed at late times, and immune effector levels do not collapse following the initial response.

Virus-specific T cells decline rapidly in all scenarios, indicating that they do not drive long-term control. Instead, separation between outcomes is dominated by CAR T persistence and the accumulation of memory. In the *Cure-tuned++* regime, the memory variable attains the highest sustained level, and CAR T cells remain stable rather than contracting, corresponding to durable tumor suppression rather than transient control followed by relapse.

### 3.6 Memory Sustains CAR T Levels Beyond the Viral Dosing Window

To isolate the mechanistic role of memory, we directly contrast CAR T dynamics in the memory-free and memory-extended formulations. Figure 6 shows the baseline model without memory, while Figure 7 displays the corresponding memory-enabled dynamics under identical treatment structures.

In the absence of memory, CAR T cells exhibit a transient expansion that is tightly coupled to viral exposure. Following the decline of viral load, CAR T levels contract steadily across all parameter regimes, even in settings that achieve strong early tumor reduction. This contraction coincides with the late expansion of antigen-negative tumor compartments, indicating that short-lived CAR T responses are insufficient for durable control.

By contrast, in the memory-extended model, CAR T trajectories diverge qualitatively. While the initial CAR T expansion remains comparable to the memory-free case, the subsequent dynamics differ fundamentally. When memory is present, CAR T levels stabilize after the viral dosing window rather than declining, with the strongest stabilization observed in the *AGG+* and *Cure-tuned++* regime. This sustained CAR T presence correlates directly with long-term suppression of antigen-negative tumor cells.

Notably, virus-specific T cells decay rapidly in both formulations and across all parameter regimes, demonstrating that memory does not act by prolonging virus-specific immunity. Instead, the distinguishing feature of durable tumor control is CAR T persistence itself. These results indicate that peak CAR T magnitude is not predictive of outcome; rather, the maintenance of CAR T levels at late times determines whether transient tumor control is converted into sustained suppression.

Together, these findings show that memory does not amplify the initial CAR T response. Instead, it stabilizes CAR T levels after viral exposure declines, providing a mechanistic link between immune recall and durable tumor control.

### 3.7 Interaction Between Memory, Viral Boosting, and Immune Exhaustion

Although PD-1/PD-L1–mediated inhibitory signaling remains active following immune engagement, memory fundamentally alters the late-time balance between immune exhaustion and effector persistence. Across all treatment settings, PD-1/PD-L1 signaling is maintained at comparable levels, indicating sustained immune regulation rather than progressive release from inhibition.

Across all parameter regimes, PD-1/PD-L1 signaling remains active throughout the response, irrespective of whether memory is present. Thus, immune exhaustion persists as a regulatory pathway in both memory-free and memory-enabled formulations.

#### Without memory

Effector contraction dominates at late times. Following the decline of viral stimulation, CAR T and tumor-specific T-cell populations progressively contract, while PD-1/PD-L1–mediated inhibitory signaling remains active. As effector levels fall below a functional threshold, the regulatory axis effectively dominates the immune response. This shift coincides with loss of tumor control, particularly through the expansion of antigen-negative tumor compartments.

#### With memory

Effector persistence dominates at late times. In memory-enabled regimes, CAR T and tumor-specific T-cell activity remain sufficient despite continued PD-1/PD-L1 signaling. Rather than eliminating immune exhaustion, memory shifts the balance between exhaustion and persistence, allowing sustained immune pressure to be maintained after viral exposure declines.

Importantly, this behavior does not rely on persistent viral boosting. Memory enables CAR T and tumor-specific T-cell function to remain stable beyond the viral dosing window, demonstrating that durable tumor control arises from immune recall rather than prolonged viral presence.

Finally, this interaction is not restricted to a single finely tuned parameter set. Durable tumor suppression in the presence of regulatory signaling emerges consistently across a class of biologically plausible memory-enabled parameter regimes, underscoring the robustness of the memory-mediated persistence mechanism.

## 4 Discussion

In this study, we developed a unified mechanistic model that integrates oncolytic virotherapy, CAR T cell dynamics, innate and adaptive immunity, and PD-1/PD-L1/PD-L1-mediated regulation within a single framework. By explicitly resolving antigen-positive and antigen-negative tumor subpopulations and coupling viral infection, immune activation, and checkpoint inhibition, the model captures several of the core biological barriers that limit CAR T efficacy in solid tumors: antigen heterogeneity, transient effector persistence, and the suppressive tumor microenvironment. This integrated structure allows us to examine both early tumor responses and the mechanisms that govern long-term control or relapse.

Consistent with the model structure described above, the strong early tumor regression followed by relapse observed in our simulations reflects a commonly reported experimental pattern in solid-tumor CAR T therapy. Clinically, IL13R*α*2-targeted CAR T therapy in recurrent multifocal glioblastoma induced a dramatic but transient complete response that was ultimately followed by disease recurrence at new anatomical sites, with evidence suggesting reduced target antigen expression [1]. Similarly, preclinical studies in orthotopic glioblastoma xenograft models demonstrate that CAR T therapies targeting a single tumor antigen often produce marked initial tumor regression, yet relapse occurs in a substantial fraction of treated animals, underscoring the impact of antigen heterogeneity and limited CAR T persistence [10]. More broadly, xenograft studies across solid tumors report that early CAR T–mediated tumor control is frequently followed by relapse driven by antigen loss, reinforcing antigen-negative escape as a central barrier to durable therapeutic efficacy [11]. Together, these findings support that the transient responses and late tumor regrowth captured by our model represent widely observed behaviors in solid-tumor CAR T systems rather than exceptional outcomes.

Global sensitivity analysis of the memory-free system revealed that a small number of dominant biological axes govern treatment outcomes. In particular, antigen-negative tumor growth and infection dynamics determine whether the system remains tumor-dominated or enters a virus-driven regression regime, while T-cell killing efficiency and survival shape the magnitude and duration of immune pressure. These findings align with prior modeling and experimental studies identifying antigen-negative escape and insufficient T-cell persistence as central obstacles to durable control in solid tumors [17]. Importantly, strengthening these axes within biologically plausible ranges improved early responses but was insufficient to prevent late tumor regrowth. Unlike previous OV–CAR T modeling studies that emphasize enhancing early cytotoxic synergy or checkpoint relief, our framework demonstrates that strengthening these mechanisms alone is insufficient to alter late-time relapse in the absence of sustained immune memory.

Consistent with this sensitivity structure, experimental efforts in solid tumors that focus on enhancing cytotoxic potency, dose intensity, or tumor infiltration have generally succeeded in amplifying early tumor regression but have not altered the propensity for relapse [1, 10, 11] These findings indicate that such strategies primarily reshape early tumor trajectories without changing the underlying late-time system behavior, explaining why dose escalation and CAR optimization approaches often plateau in efficacy. Within the context of our model, this limitation arises because strengthening early effector mechanisms does not address the decay of immune pressure or the unchecked expansion of antigen-negative tumor populations at later times.

Motivated by experimental evidence that oncolytic viruses can expand dual-specific CAR T populations and induce memory-like phenotypes, we extended the model to include a reduced representation of virus-driven immune memory. This extension does not rely on detailed phenotypic subdivision but instead captures the essential functional consequences of memory: persistence beyond the initial response and enhanced recall upon subsequent viral exposure. Within this framework, memory fundamentally alters system-level behavior. While memory-free scenarios consistently relapse, memory-enabled regimes sustain CAR T levels, maintain immune pressure at later time points, and suppress antigen-negative tumor growth over extended periods.

This behavior is also consistent with experimental observations from oncolytic virus–CAR T combination studies, providing further biological grounding for the memory-enabled regime of the model. In particular, Evgin et al.[8] demonstrated that oncolytic virotherapy can promote sustained CAR T persistence and prolonged tumor control by expanding functionally durable, dual-specific T cell populations. In those experiments, CAR T cells remained detectable and functionally active at late time points and supported extended immune pressure on the tumor, even without explicit resolution of classical memory subsets. In qualitative agreement with these findings, the memory-enabled regime in our model sustains CAR T levels beyond the initial response phase, maintains immune pressure at late times, and suppresses antigen-negative tumor growth over extended periods. Notably, this agreement emerges despite the use of a reduced representation of immune memory that captures functional persistence and recall rather than detailed phenotypic subdivision. Together, these results suggest that the system-level consequences of virus-induced immune memory—rather than the specific memory phenotypes themselves—are sufficient to fundamentally alter long-term tumor control dynamics.

A key insight from the memory-extended simulations is that durable tumor control arises not from amplifying early cytotoxicity, but from stabilizing immune activity after viral and antigenic signals wane. In the model, memory primarily maintains CAR T responsiveness rather than increasing peak expansion. This distinction helps reconcile why aggressive early interventions can fail while strategies that promote immune recall succeed. It also reframes the role of oncolytic viruses in combination therapy: beyond direct oncolysis or transient inflammation, viral exposure functions as an immune programming signal that sustains long-term antitumor immunity.

Beyond static measures of sustained CAR T persistence, experimental studies also report recall-type responses following secondary stimulation, further supporting a memory-based mechanism of durable tumor control. In particular, Evgin et al.[8] demonstrated that a second in vivo viral boost can actively restimulate memory CAR T cells and restore antitumor efficacy. In their model, an initial treatment with virus-loaded CAR T cells was insufficient to achieve long-term tumor control; however, subsequent intravenous re-dosing with VSV-mIFN*β* triggered a robust recall response, leading to durable tumor protection in the majority of treated animals. This effect was antigen-specific, as boosting with an unrelated virus failed to restimulate CAR T activity, highlighting the role of virus-specific immune memory rather than nonspecific inflammation. Consistent with these findings, the memory-enabled regime in our model exhibits enhanced responsiveness upon renewed viral or antigenic stimulation, allowing immune pressure to be re-established after the initial response declines. In this way, the model provides a mechanistic explanation for how viral re-dosing or antigen re-exposure can rescue antitumor immunity through recall dynamics, even without explicitly resolving individual memory T cell subsets.

The PD-1/PD-L1 axis emerges in this framework as a modulator rather than a sole determinant of long-term efficacy. While immune activation initially elevates inhibitory signaling, memory-enabled persistence allows effector populations to overcome exhaustion and maintain tumor control. This suggests that checkpoint blockade alone may be insufficient without mechanisms that preserve immune competence over time, but that it may synergize strongly with strategies that enhance memory or recall.

Building on these observations, several preclinical studies combining oncolytic virotherapy with CAR T cells and immune checkpoint blockade further support the idea that durable therapeutic success depends on immune persistence rather than transient relief of T cell exhaustion alone. For example, an oncolytic adenovirus engineered to deliver a bispecific T cell engager, IL–12, and a PD-L1-blocking antibody markedly enhanced CAR T–mediated tumor control in heterogeneous solid tumor models, yet long-term efficacy varied across settings [25]. Similarly, adenovirotherapy delivering IL–12 together with PD–L1 blockade augmented CAR T activity and prevented loss of CAR T cells at the tumor site, indicating enhanced immune persistence beyond checkpoint inhibition alone [28]. In related work, cytokine-armed oncolytic adenoviruses combined with CAR T cells increased tumor infiltration and prolonged T cell function, even in the absence of explicit checkpoint blockade, further implicating sustained immune activity as a key determinant of outcome [35]. Together, these studies suggest that while checkpoint inhibition can alleviate exhaustion, durable tumor control emerges primarily when combination strategies promote prolonged CAR T persistence and functional recall. Consistent with this interpretation, our model reproduces these qualitative behaviors by demonstrating that long-term control arises from stabilized immune activity rather than amplified early cytotoxicity alone.

Our model reproduces experimentally observed behaviors across broad parameter regimes, underscoring the robustness of the underlying mechanisms. At the same time, several limitations remain. The model is spatially lumped, does not resolve trafficking or spatial heterogeneity, and is not calibrated to a specific experimental dataset. The memory model is intentionally simplified, aggregating several biological processes, and toxicity or optimal scheduling is not addressed. Nonetheless, the qualitative behaviors identified here are robust across parameter regimes and consistent with experimental observations, and our choices make the model broadly applicable across tumor types, viral platforms, and CAR constructs. By collapsing complex biological processes into a small number of interpretable control axes, the framework provides a tractable platform for hypothesis generation and experimental design.

A practical implication of the framework is that it can help prioritize OV designs by distinguishing mechanisms that primarily amplify *early inflammation* from those that promote *long-term CAR T persistence and recall*. In the model, interventions that increase acute innate activation (e.g., transient cytokine spikes) can deepen early tumor regression, yet they often leave the late-time attractor unchanged if CAR T levels still decay rapidly and antigen-negative growth remains unchecked. By contrast, OV modifications that effectively reduce the post-peak contraction of CAR T cells or enable recall upon renewed viral-antigenic exposure shift the system toward sustained immune pressure, delaying or preventing late regrowth. Experimentally, this suggests prioritizing OV candidates based not only on early tumor shrinkage, but on late-time readouts such as (i) persistence of CAR T cells at the tumor site beyond the initial response window, (ii) maintenance of functional markers and cytotoxic capacity at later time points, and (iii) the magnitude of recall responses after viral re-dosing or antigen re-stimulation. In this way, the model provides a mechanistic rationale for why OV platforms that program durable persistence can outperform approaches that merely intensify early inflammation.

This level of abstraction allows general conclusions to be drawn across tumor types and therapeutic designs, rather than tailoring predictions to a single experimental system. At the same time, this abstraction necessarily omits several biologically important mechanisms that may influence therapeutic outcomes in specific experimental settings. Future work will focus on extending this framework along several complementary directions to further strengthen its experimental relevance and predictive utility. One important next step is calibration of the model to specific preclinical datasets. This would enable quantitative testing of whether memory-driven persistence alone can explain observed outcomes, or whether additional mechanisms—such as spatial heterogeneity, trafficking limitations, or cytokine-mediated toxicity—must be incorporated. Incorporating spatial structure, either through partial differential equations or coupling to agent-based models, would allow investigation of how localized viral infection, heterogeneous antigen expression, and immune infiltration patterns shape long-term control. In parallel, the memory representation introduced here could be refined to distinguish between effector, central memory, and stem-like CAR T phenotypes, informed by emerging single-cell and lineage-tracing data. Such extensions would allow direct comparison between different CAR designs and viral constructs that preferentially promote specific memory programs. Finally, the framework could be leveraged to explore treatment optimization questions, including the timing and magnitude of viral re-dosing, coordination with CAR T infusion schedules, and combination with checkpoint blockade. By systematically interrogating these design choices in silico, the model can help prioritize experimentally testable strategies for converting early responses into durable remissions. Overall, this work provides a unified, mechanistic explanation for why early success in OV–CAR T combination therapy often fails to translate into durable remission, and why memory induction may be a critical missing ingredient. By linking viral dynamics, immune regulation, and CAR T persistence within a single framework, the model offers both conceptual clarity and practical utility. It supports the view that converting transient responses into lasting control will require therapies designed not only to kill tumor cells, but also to program long-term immune competence. Our model and proposed extension can serve as a tractable platform for interpreting experimental outcomes, generating testable hypotheses, and guiding strategies to transform transient responses into lasting remission in solid tumors.

## Declaration of Competing Interests

The authors declare that they have no competing interests.

## Funding

This research did not receive any specific grant from funding agencies in the public, commercial, or not-for-profit sectors.

## Appendix A.

**Memory-Free Parameterization**

To identify the dominant parameters governing the coupled OV–CAR T–immune dynamics, we performed a global sensitivity analysis using Partial Rank Correlation Coefficients (PRCC). The analysis was conducted on Latin Hypercube samples spanning the full parameter ranges and evaluated at four representative time points (*t*=15, 30, 60, and final).

The resulting sensitivity patterns revealed the parameters most strongly shaping tumor and viral kinetics—particularly *r*_*T N*_, *K*_*N*_, *β*_*NV*_, and *δ*_*NI*_ on the tumor–infection axis, and *k*_*T A*_, *h*_*V*_, and *δ*_*Y T*_ on the immune axis. Guided by these PRCC rankings, two parameter sets were then derived as shown in Table 3: a baseline aggressive (AGG) configuration and an enhanced activation (AGG+) configuration, each emphasizing distinct regimes of infection and immune response. Although the viral burst size *b*_*T*_ and viral decay rate *ω* do not rank among the top PRCC-sensitive parameters, they enter the viral-load equation as the dominant production and clearance terms. Plausible variations in *b*_*T*_ and *ω* therefore primarily rescale the magnitude and duration of viral exposure without introducing a new control axis. Accordingly, modest adjustments to *b*_*T*_ and *ω* are included in both AGG and AGG+ to reflect uncertainty in viral production and decay while remaining consistent with the PRCC-guided structure.

**Table 3:**
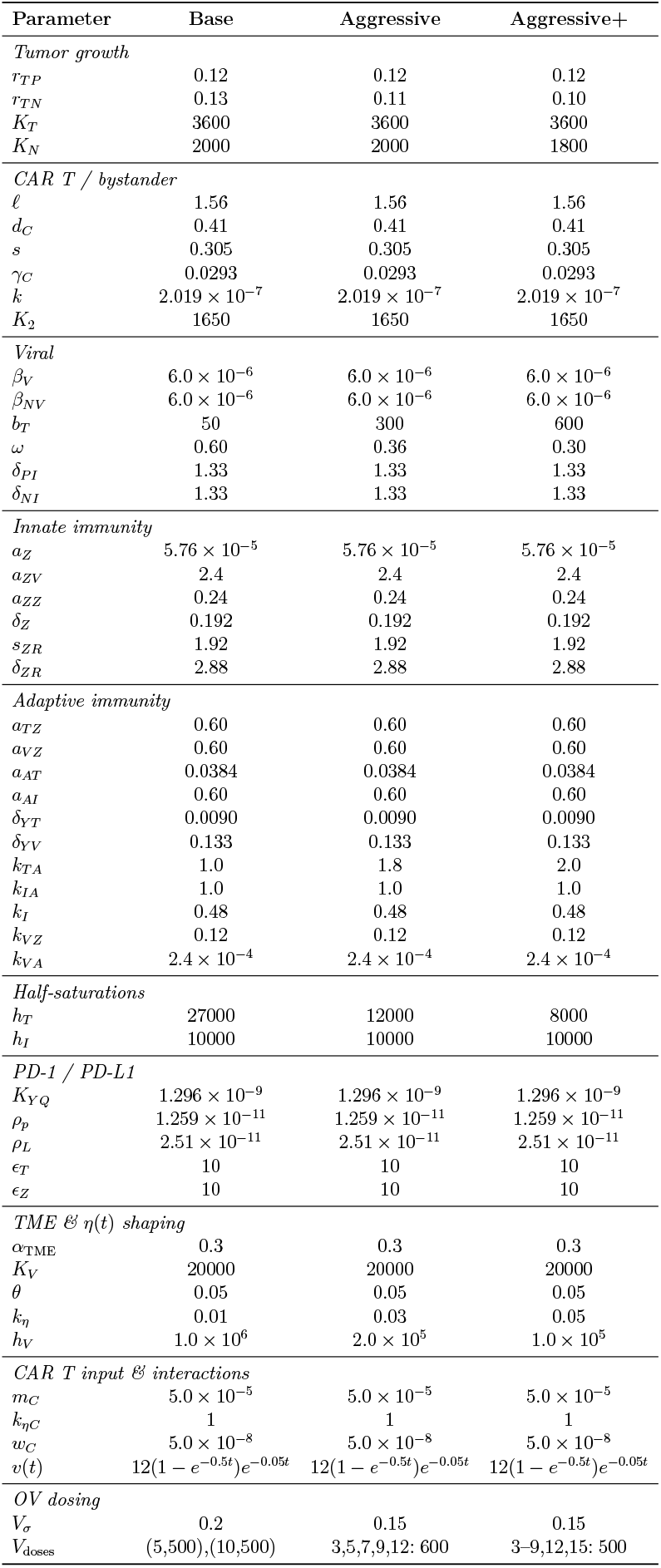
Parameter sets used in simulations (guided by [30, 17]).

## Appendix B.

**Memory-Extension Parameterization**

### B.1 Biological Motivation and Rationale

Recent experimental work has demonstrated that oncolytic viruses (OVs) can prime CAR T cells through stimulation of the native TCR with viral or virally encoded epitopes, resulting in enhanced proliferation, broadened antitumor function, and the formation of durable memory phenotypes. These findings, reported in Evgin et al. [8], motivate the introduction of a memory variable *M* (*t*) in our extended CART+OV model.

The memory induction rate was chosen as *k*_mem_ = 0.15 − 0.30 day^−1^ so that OV exposure yields a measurable memory response within 3–7 days, consistent with the expansion kinetics of dual-specific (DS) CAR T cells. The half-saturation parameter *h*_mem_ was set to reflect that memory induction is driven by intermediate-to-high viral loads, as suggested by the viral boosting results of [8]. The decay rate *δ*_mem_ = 0.004−0.01 day^−1^ produces a long-lived memory pool with an effective half-life of 70–170 days, in line with the observation that DS CAR T cells remain detectable in the blood and spleen for over 100 days following adoptive transfer.

Functional enhancement is represented by the multiplicative factor *α*_mem_ = 0.8−1.4, which increases the effective CAR-mediated killing rate by approximately 1.8–2.5-fold when memory is present. This range is consistent with the general observation in [8] that OV-driven TCR stimulation can strengthen overall CAR T performance.A small additive term *σ*_*M*_ = 0.02 introduces a minor numerical contribution to the CAR T related terms whenever memory is present. The drift parameter *k*_spread_ = 0.005 day^−1^ encodes a slow redistribution between the virus-responsive (*Y*_*V*_) and tumor-responsive (*Y*_*T*_) T–cell compartments. Both mechanisms are incorporated in a phenomenological manner and are consistent with the general observation in [8] that OV-driven TCR stimulation can broaden the overall response of CAR T cell populations.

### B.2 Memory-Related Parameters

The parameter values in Appendices A and B.2 were chosen to stay within ranges reported in earlier modelling studies rather than to fit a specific dataset exactly. Tumor growth rates, carrying capacities, and immune killing rates were taken in the same order of magnitude as those used by Storey et al. [30] and Kara et al. [17], and then adjusted to match our choice of state variables measured in volume (mm^3^) instead of cell counts. Viral infection, production, and clearance parameters were similarly selected so that the resulting time scales and amplitudes are consistent with oncolytic virus models in the literature. The memory-related parameters were chosen so that the build-up and decay of *M* (*t*) occur on time scales comparable to the expansion and persistence of dual-specific CAR T cells described by Evgin et al. [8], while keeping their effect modest. Overall, the Base, Aggressive, Aggressive+, and Cure-tuned++ sets, as shown in Table 4, are intended to represent biologically plausible scenarios that illustrate the qualitative behaviour of the combined OV–CAR T–immune system.

**Table 4:** Parameter sets used in simulations (guided by [8, 30, 17]).

| Parameter | Base | Aggressive | Aggressive+ | Cure-tuned++ |
| --- | --- | --- | --- | --- |
| $r_{TP}$ | 0.12 | 0.12 | 0.12 | 0.12 |
| $r_{TN}$ | 0.13 | 0.11 | 0.10 | 0.10 |
| $K_T$ | 3600 | 3600 | 3600 | 3600 |
| $K_N$ | 2000 | 2000 | 1800 | 1800 |
| $\ell$ | 1.56 | 1.56 | 1.56 | 1.56 |
| $d_C$ | 0.41 | 0.41 | 0.41 | 0.41 |
| $s$ | 0.305 | 0.305 | 0.305 | 0.305 |
| $\gamma_C$ | 0.0293 | 0.0293 | 0.0293 | 0.0293 |
| $k$ | $2.019 \times 10^{-7}$ | $2.019 \times 10^{-7}$ | $2.019 \times 10^{-7}$ | $2.019 \times 10^{-7}$ |
| $K_2$ | 1650 | 1650 | 1650 | 1650 |
| $\beta_V$ | $6.0 \times 10^{-6}$ | $6.0 \times 10^{-6}$ | $6.0 \times 10^{-6}$ | $6.0 \times 10^{-6}$ |
| $\beta_{NV}$ | $6.0 \times 10^{-6}$ | $9.6 \times 10^{-6}$ | $1.2 \times 10^{-5}$ | $1.68 \times 10^{-5}$ |
| $b_T$ | 50 | 300 | 600 | 700 |
| $\omega$ | 0.60 | 0.36 | 0.30 | 0.288 |
| $\delta_{PI}$ | 1.33 | 1.33 | 1.33 | 1.33 |
| $\delta_{NI}$ | 1.33 | 1.33 | 1.33 | 1.33 |
| $a_Z$ | $5.76 \times 10^{-5}$ | $5.76 \times 10^{-5}$ | $5.76 \times 10^{-5}$ | $5.76 \times 10^{-5}$ |
| $a_{ZV}$ | 2.4 | 2.4 | 2.4 | 1.92 |
| $a_{ZZ}$ | 0.24 | 0.24 | 0.24 | 0.24 |
| $\delta_Z$ | 0.192 | 0.192 | 0.192 | 0.192 |
| $s_{ZR}$ | 1.92 | 1.92 | 1.92 | 1.92 |
| $\delta_{ZR}$ | 2.88 | 2.88 | 2.88 | 2.88 |
| $a_{TZ}$ | 0.60 | 0.60 | 0.60 | 0.60 |
| $a_{VZ}$ | 0.60 | 0.60 | 0.60 | 1.92 |
| $a_{AT}$ | 0.0384 | 0.0384 | 0.0384 | 0.0384 |
| $a_{AI}$ | 0.60 | 0.60 | 0.60 | 1.56 |
| $\delta_{YT}$ | $9.0 \times 10^{-4}$ | $9.0 \times 10^{-4}$ | $9.0 \times 10^{-4}$ | $9.0 \times 10^{-4}$ |
| $\delta_{YV}$ | 0.133 | 0.133 | 0.133 | 0.133 |
| $k_{TA}$ | 1.0 | 1.8 | 2.0 | 2.0 |
| $k_{IA}$ | 1.0 | 1.0 | 1.0 | 2.8 |
| $k_I$ | 0.48 | 0.48 | 0.48 | 1.68 |
| $k_{VZ}$ | 0.12 | 0.12 | 0.12 | 0.12 |
| $k_{VA}$ | $2.4 \times 10^{-4}$ | $2.4 \times 10^{-4}$ | $2.4 \times 10^{-4}$ | $2.4 \times 10^{-4}$ |
| $h_T$ | 27000 | 12000 | 8000 | 8000 |
| $h_I$ | 10000 | 10000 | 10000 | 2500 |
| $K_{YQ}$ | $1.296 \times 10^{-9}$ | $1.296 \times 10^{-9}$ | $1.296 \times 10^{-9}$ | $1.296 \times 10^{-9}$ |
| $\rho_p$ | $1.259 \times 10^{-11}$ | $1.259 \times 10^{-11}$ | $1.259 \times 10^{-11}$ | $1.259 \times 10^{-11}$ |
| $\rho_L$ | $2.51 \times 10^{-11}$ | $2.51 \times 10^{-11}$ | $2.51 \times 10^{-11}$ | $2.51 \times 10^{-11}$ |
| $\epsilon_T$ | 10 | 10 | 10 | 10 |
| $\epsilon_Z$ | 10 | 10 | 10 | 10 |
| $\alpha_{TME}$ | 0.3 | 0.3 | 0.3 | 0.3 |
| $K_V$ | 20000 | 20000 | 20000 | 20000 |
| $\theta$ | 0.05 | 0.05 | 0.05 | 0.08 |
| $k_\eta$ | 0.01 | 0.03 | 0.05 | 0.02 |
| $h_V$ | $1.0 \times 10^6$ | $2.0 \times 10^5$ | $1.0 \times 10^5$ | $5.0 \times 10^5$ |
| $k_{mem}$ | 0.15 | 0.15 | 0.15 | 0.30 |
| $h_{mem}$ | 200 | 200 | 200 | 280 |
| $\delta_{mem}$ | 0.01 | 0.01 | 0.01 | 0.004 |
| $\alpha_{mem}$ | 0.8 | 0.8 | 0.8 | 1.4 |
| $\sigma_M$ | 0.02 | 0.02 | 0.02 | 0.02 |
| $k_{spread}$ | 0.005 | 0.005 | 0.005 | 0.005 |
| $m_C$ | $5.0 \times 10^{-5}$ | $5.0 \times 10^{-5}$ | $5.0 \times 10^{-5}$ | $4.0 \times 10^{-5}$ |
| $k_{\eta C}$ | 1 | 1 | 1 | 1 |
| $w_C$ | $5.0 \times 10^{-8}$ | $5.0 \times 10^{-8}$ | $5.0 \times 10^{-8}$ | $4.0 \times 10^{-8}$ |
| $V_\sigma$ | 0.2 | 0.15 | 0.15 | 0.15 |
| $V_{doses}$ | (5,500),(10,500) | 3-9,12,15:600 | 3-9,12,15:700 | 3-9,12,15,18,21,24,28:700 |

## Notes

### Competing Interest Statement

The authors have declared no competing interest.

